# Engineered Peptide-Functionalized Hydrogels Modulate the RNA Transcriptome of Human Nucleus Pulposus Cells In Vitro

**DOI:** 10.1101/2021.03.05.434094

**Authors:** Marcos N. Barcellona, Julie E. Speer, Liufang Jing, Munish C. Gupta, Jacob M. Buchowski, Michael P. Kelly, Lori A. Setton

## Abstract

Degeneration and aging of the nucleus pulposus (NP) of the intervertebral disc (IVD) is accompanied by alterations in NP cell phenotype marked by a shift towards a fibroblast-like, catabolic state. We have recently demonstrated an ability to manipulate the phenotype of human adult degenerative NP cells through 2D culture upon poly(ethylene glycol) (PEG) based hydrogels dually functionalized with integrin- and syndecan-binding laminin-mimetic peptides (LMPs). In the present study, we sought to understand the transcriptomic changes elicited through NP cell interactions with the LMP-functionalized hydrogel system (LMP gel) by examining targets of interest *a priori* and by conducting unbiased analysis to identify novel mechanosensitive targets. The results of gene specific analysis demonstrated that the LMP gel promoted adult degenerative NP cells to upregulate 148 genes including several NP markers (e.g. NOG and ITGA6) and downregulate 277 genes, namely several known fibroblastic markers. Additionally, 13 genes associated with G protein-coupled receptors, many of which are known drug targets, were identified as differentially regulated following culture upon the gel. Furthermore, through gene set enrichment analysis we identified over 700 pathways enriched amongst the up- and downregulated genes including pathways related to cell differentiation, notochord morphogenesis, and intracellular signaling. Together these findings demonstrate the global mechanobiological effects induced by the LMP gel and confirm the ability of this substrate to modulate NP cell phenotype.

## 1. Introduction

The nucleus pulposus (NP) of the intervertebral disc (IVD) has been observed to undergo considerable changes in matrix composition and overall biochemical makeup with age and maturation ^1–5^. The cells of the NP are derived from the embryonic notochord, a structure which expresses an array of signaling molecules and transcription factors such as brachyury (T), noggin (NOG), and notochord homeobox (NOTO), some of which are observed to disappear during early developmental stages ^4,6^. Both notochordal and juvenile NP cells secrete a matrix rich in collagens, predominantly collagen type II, proteoglycans, and other proteins such as laminins and fibronectin ^2,7,8^.

With degeneration, there is a loss of hydration in the NP of the IVD that contributes to tissue stiffening and reduced IVD height, with NP shear moduli values increasing from <1 kPa in healthy NP to 10-20 kPa in moderately degenerated NP ^1,7,9–11^. These changes are associated with a decrease in NP cellularity and a shift towards more fibroblast-like cell phenotype with altered biosynthesis including secretion of a matrix with decreased sulfated glycosaminoglycans (sGAGs), decreased collagen type II, and increased collagen type I ^12–14^. Changes have also been observed in the presence of matrix degrading enzymes such as the MMP and ADAMTS families, which results in an altered cellular microenvironment and a decreased ability for tissue repair and homeostasis ^4,15–17^. As the NP has limited ability for self-repair, biomaterials have been investigated as a means to deliver cells to the degenerative disc or to provide stimuli to modulate cell (endogenous or exogenous) behavior and promote IVD regeneration ^14,18,19^.

The mechanobiological cues cells encounter in their environment have the ability to regulate cell behaviors (including cell signaling, biosynthesis, and motility), phenotype, and global transcriptomic changes ^20–23^. Biomaterial strategies have utilized natural and/or synthetic materials functionalized with bioactive moieties such as full-length proteins or peptide sequences in order to promote tissue repair or cell differentiation ^19,24–26^. In addition to ligand presentation, other biomaterial parameters including substrate stiffness, porosity, topography, and surface chemistry have been tested for their ability to modulate cell behaviors ^27–33^.

^14,20,34–37^

Previous studies have shown that culture of NP cells upon soft (∼0.5 kPa) hydrogel substrates functionalized with laminin-111 can be used to promote the cell behaviors and gene expression profiles characteristically observed in the juvenile NP, while cells cultured on stiff (10-20 kPa) hydrogel substrates or tissue culture plastic exhibit a fibroblast-like phenotype ^9,38,39^. While this work made use of full-length laminin proteins, short peptide sequences offers several advantages in biomaterial design including the opportunity to precisely control sites of biomaterial and cell interactions ^24,39^. A screen of 10 laminin-mimetic peptides (LMPs) demonstrated that LMPs including the syndecan-binding peptide AG73 or integrin-binding peptide IKVAV promoted NP cell attachment at levels that were comparable to full length Laminin-111 ^39^. As syndecans and integrins can work synergistically to regulate cell adhesion and intracellular signaling ^40,41^, we have recently developed poly(ethylene glycol) (PEG) gels which co-presented the LMPs IKVAV and AG73 in equal molar ratios ^14^. Using this biomaterial system, we have demonstrated that stiff (15% PEG) gels functionalized with a total peptide density of 100 µM could lead to a partial re-expression of a panel of juvenile markers for human degenerate NP cells in a manner that replicated the effects of soft laminin presenting materials ^14^. Stiff biomaterials may have improved resilience under mechanical loading in the IVD space and improved injectability. Thus, the development of a stiff, LMP-functionalized gel that replicates the effects of softer substrates represents a clinically-relevant biomaterial. In the present study we sought to quantify the global transcriptomic changes and pathways activated by the LMP gel in order to both explore pathways of *a priori* interest and to use unbiased analysis to identify novel targets of the biomaterial-induced mechanotransduction.

## 2. Materials and Methods

### 2.1. NP Cell Isolation

Primary adult human NP cells were isolated from to-be-discarded surgical tissue of patients with degeneration-associated complications (n ≥ 3 for all studies, both sexes, ages 18-65, in accordance with procedures from Washington University Institutional Review Board) as previously described^39^. The NP cells were expanded on tissue culture polystyrene (TCPS) in Ham’s F12 media (Thermo Fisher Scientific, Waltham, MA) supplemented with 10% FBS and 1% penicillin-streptomycin (21% O_2_, 5% CO_2_); cells for this experimentation were used no later than passage 4.

### 2.2 Hydrogel Preparation

PEG hydrogels functionalized with LMPs (LMP gels) were prepared as previously described ^14^. Briefly, lyophilized, cysteine terminated IKVAV and AG73 peptides (full sequences for IKVAV and AG73: CSRARKQAASIKVAVSADR, and CGGRKRLQVQLSIRT respectively, GenScript Biotech, Piscataway, NJ) were dissolved in 1X PBS pH 3.25. Peptide solutions were added to a maleimide terminated 8-arm star poly(ethylene glycol) (PEG-8MAL, MW 20K, Creative PEGWorks, Durham, NC) for the functionalization of the 8-arm PEG backbone ^42^. A small PEG-dithiol (SH-PEG-SH, MW 600, Creative PEGWorks) was likewise dissolved in 1X PBS pH 3.25, and then added to the peptide-PEG-8MAL solution to initiate crosslinking and hydrogel formation. The gel solution was pipetted into 4-well chamber slides (Nunc Lab-Tek Chamber Slide Systems ™, ThermoFisher Scientific) and gels were permitted to crosslink for 1 hour, then neutralized with 1X PBS pH 7.4 and allowed to swell to equilibrium volume. LMP gels were made at 15% PEG w/v, previously measured to have a stiffness ∼10 kPa, and functionalized with 100 µM total peptide using equimolar amounts of IKVAV and AG73 (50µM of each peptide) ^14^.

### 2.3 RNA Sequencing

Primary adult human NP cells from each of three patients (24-year-old female, 53-year-old female, and 55-year-old male) were seeded in triplicate upon LMP gels or uncoated tissue culture polystyrene (TCPS) at a density of 100,000 cells per well in a 4-well chamber slide, consistent with previously published protocols ^14,43^. Cells were cultured for 4 d at 37°C and 5% CO_2_. Following the incubation period, the cells were lysed using RLT buffer (Qiagen, Hilden, Germany) + 1% mercaptoethanol and stored at - 80°C until ready for RNA isolation. Isolation was achieved using the Qiagen Mini Kit following the manufacturer’s instructions (Qiagen, Hilden, Germany). RNA qualities were measured using a Bioanalyzer (Agilent, Santa Clara, CA), and all samples used had an RNA integrity number (RIN) > 9.

RNA sequencing was performed by the Genome Technology Access Center (GTAC) at Washington University in St. Louis. Briefly, ds-cDNA was made using Clontech SMARTer® Ultra Low RNA Kit (Clontech, Mountain View, CA) following manufacturer’s recommendations. cDNA was fragmented using a Covaris E220 sonicator (peak incident power = 18, duty factor = 20%, cycles per burst = 50, for 120 seconds). Ligated fragments were amplified for 12 cycles using primers incorporating dual index tags. Fragments were then run on an Illumina NovaSeq (San Diego, CA) reading 150 bases from both ends to a depth of 30 million reads. FASTQ files provided by GTAC were used in subsequent analysis.

Using Partek Flow® software (Partek Inc., St. Louis, MO), unaligned reads were trimmed prior to alignment to the human genome (STAR 2.6.1d and referencing to the human genome hg19). Gene Specific Analysis (GSA) was conducted in Partek Flow® and used to perform downstream analysis (identification of differentially expressed genes, principal component analysis, hierarchical clustering, and gene set enrichment) using methodology as previously described ^43^. GSA was used to identify a list of upregulated and downregulated genes. First, the GSA was used to calculate fold change values (expression on LMP gel relative to expression on TCPS) and a LIMMA algorithm ^44^ was applied to conduct pair-wise comparisons of gene expression data between samples on each substrate with a calculation of a *p*-value for each gene. A gene was considered to be differentially regulated if the fold change value was greater than or equal to 2 (upregulated) or less than or equal to −2 (downregulated) and if the *p*-value was less than 0.05.

The list of differentially regulated genes was data mined to quantify gene expression for known markers of NP cells and fibrotic phenotypes (*a priori* targets) but was also used to perform unbiased analysis to identify novel targets. Furthermore, the gene lists were used to perform gene set enrichment (referring to Gene Ontology, GO, terms per the 2017_11_01 Gene Set Database) to provide biological context to the lists of differentially regulated genes and to identify cellular functions and pathways that may be associated with peptide-induced mechanotransduction. The results of the gene set enrichment were also data mined to identify pathways known to be implicated in NP and IVD biology, and also to identify unprecedented pathways. The RNA sequencing dataset generated during this study was deposited in the NCBI Gene Expression Omnibus (GEO) repository (GSE154044).

### 2.4 Statistical Analysis for RNA Sequencing

All statistical processes described below were performed in Partek Flow®. Fold change and *p*-values were calculated for each gene using a GSA algorithm. As previously described, genes were considered differentially regulated at a fold change value (LMP gel relative to TCPS) ≥ 2 (upregulated) or ≤ −2 (downregulated) and a *p*-value of < 0.05. PCA analysis was performed to reduce the dimensionality of the data and a scatterplot showing the data in principle components (PC) 1-3 was generated to visualize groupings of the data. Hierarchical clustering was also performed to conduct unsupervised clustering of the samples (*n* = 3 patients) and features (genes) using the average cluster distance metric of average linkage and a point distance metric of Euclidean distance. A heat map was created to visualize these data and genes were plotted as rows and patient samples and substrate conditions were shown on columns. Normalization was performed to generate z-scores (a number that describes how many standard deviations a value is from the mean) for each gene in order to visualize patterns in gene expression by substrate and patient. Dendrograms were applied to the heat map to demonstrate the results of the hierarchical clustering. The most up- and down-regulated genes were identified as those with the highest or lowest fold change values, *p*-values < 0.05, and with known molecular function and/or biologic process gene ontology terms (UniProt database).

Gene set enrichment (GSE) was further conducted on the lists of upregulated and downregulated genes separately to identify gene sets (GO terms) significantly overrepresented in the list. To determine enrichment, the software first developed a contingency table containing 4 categories – number of genes in the list (i.e. upregulated or downregulated genes) and in the given gene set (GO term), number of genes that are not in the list but in the set, number of genes in the list but not in the set, and the number of genes not in the list and not in the set. In order to determine if a GO term was enriched in the data set, a Fisher’s exact test was used to calculate a *p*-value and an enrichment score (negative natural logarithm of the *p*-value). A higher enrichment score (and concomitantly lower *p*-values) indicate gene sets that are overrepresented by the genes in the up- or downregulated lists respectively.

### 2.5 Validation Using Immunocytochemistry

Validation of RNA sequencing results were confirmed at the protein level for select markers (n = 4; cells from both male and female patients, ages 18-65). As G protein-coupled receptors (GPCR) are commonly druggable targets, protein expression was confirmed for one upregulated and one downregulated GPCR-associated marker (Alpha-2C adrenergic receptor and Oxytocin receptor, respectively) as identified through GSA and identification of GO terms. Adult degenerative human NP cells were seeded on TCPS or LMP gels for 4 d (10,000 cells per well). Afterwards, the cells were fixed using 4% paraformaldehyde (10 min), permeabilized using Triton-X (0.2% Triton-X in PBS^+/+^; 10 min), and incubated for 30 min in blocking buffer (3.75% BSA and 5% goat serum, Thermo Fisher Scientific). Samples were then immunolabeled for Alpha-2C adrenergic receptor, also known as α2C adrenoceptor, (ADRA2C ; 1:100; Abcam, Cambridge, UK) or Oxytocin receptor (OXTR; 1:100; Proteintech, Rosemont, IL) or stained with the respective isotype control. AlexaFluor™ secondary antibodies were used to visualize the samples (Invitrogen, Carlsbad, CA, 1:200) and DAPI (2µg/mL, Sigma Aldrich, St. Louis, MO) was used to counterstain nuclei. Samples were imaged using confocal microscopy (TCS-SPE with DM6 RGBV confocal microscope; Leica DFC7000 T camera; using Leica LAS X core software; Leica Microsystems; Buffalo Grove, IL) and a 20x objective. A minimum of 5 regions of interest (ROIs) were visualized for each substrate (TCPS or LMP gel) and patient (*n* = 4). Fiji software ^45^ was utilized to quantify the expression of each protein and used to determine the mean fluorescence intensity (MFI) for each cell/cell cluster. Relative protein expression was calculated as the MFI normalized to the average MFI for cells treated with the isotype control antibody. One-tailed *t*-tests were used to compare the expression of ADRA2C and OXTR between the TCPS and LMP gel conditions.

## 3. Results

### 3.1 Global Gene Expression of NP Cells is Altered by Culture on LMP Gels Functionalized with IKVAV and AG73

PCA analysis revealed that 91.5% of the total data variance could be explained by PCs 1-3, with 50.8% explained by PC1 alone (Fig. 1a). Visualizing the PCA data as a scatterplot, it can be seen that while there was some separation of data due to inter-patient variability, gene expression in cells cultured on LMP gels clustered distinctly from cells cultured on TCPS (clusters are outlined as shown in Fig. 1a). Hierarchical clustering further confirmed that the data clustered amongst substrate condition (TCPS or LMP gel) and that the patterns of gene expression were different between the two cell culture environments. (Fig. 1b). Compared to cells on TCPS, culture on LMP gels resulted in 425 differentially regulated genes – 148 upregulated (fold change ≥ 2 and *p* < 0.05) and 277 downregulated genes (fold change ≤ −2 and *p* < 0.05; Fig. 1c).

**Figure 1.**
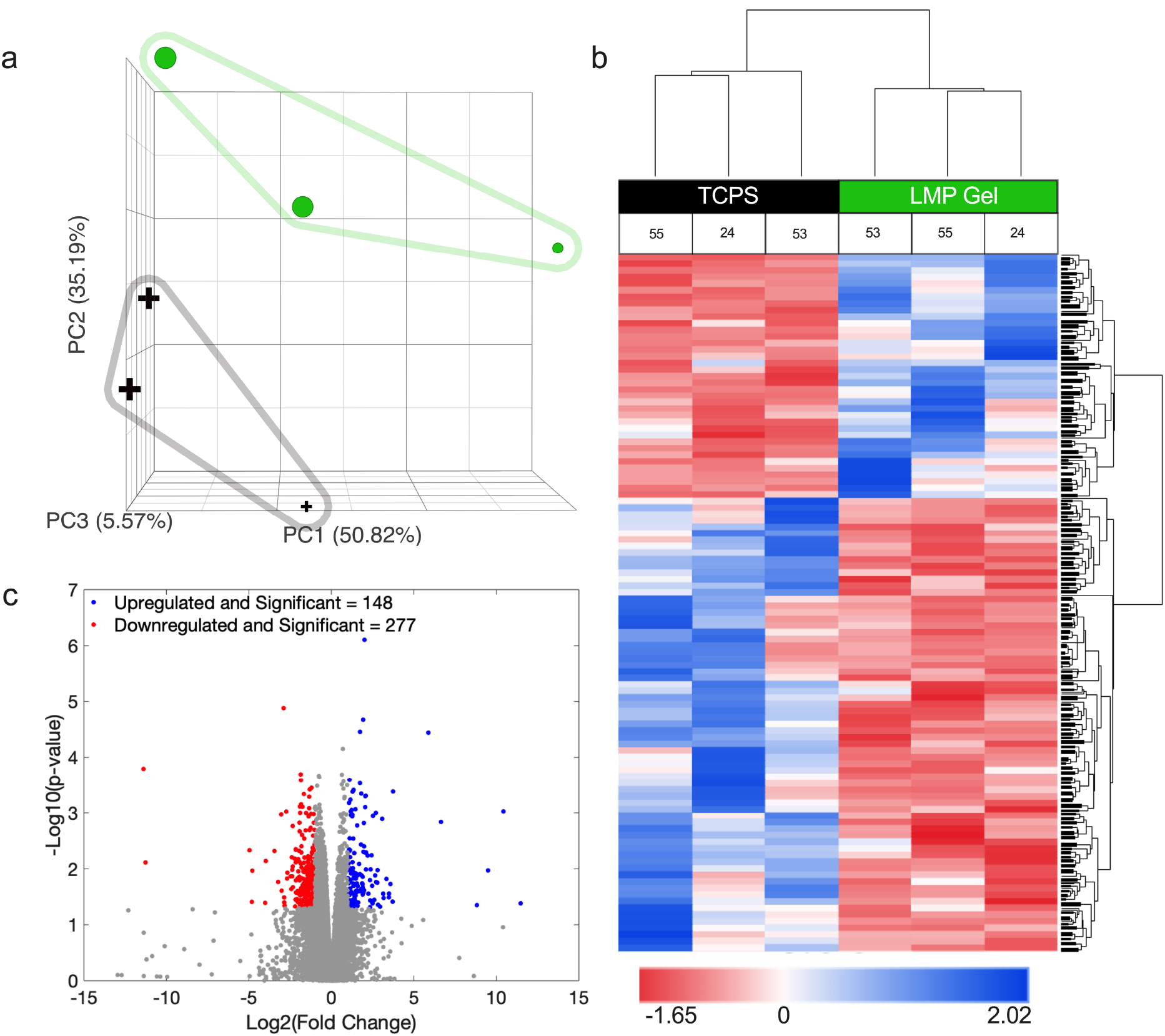
Culture of NP cells on AG73+IKVAV gels modulates global transcriptome. (a) PCA analysis of cells cultured on LMP gels (green circles) or uncoated tissue culture polystyrene (TCPS; black crosses). Each symbol represents NP cells from a human patient (24, 53, 55 –year-old, size of symbol increases with patient age). The percentage of variance explained by each PC is listed in parentheses. (b) Hierarchical clustering and heat map of data from cells on the TCPS or LMP gel. Each column shows data from a given patient as indicated by patient age (55, 24, or 53), z −score of gene expression is plotted across rows (low expression = red, high expression = blue). (c) Volcano plot shows differentially regulated genes (downregulated = red, upregulated = blue) and genes which are not differentially regulated (grey).

The most upregulated genes (with significant *p*-values) are given in Fig. 2a and Table 1 and include noggin (NOG), an important protein involved in regulating musculoskeletal system development. Many of the most upregulated genes were associated with cellular signaling and ion transport including Somatostatin (SST), Proprotein convertase subtilisin/kexin type 9 (PCSK9), and Solute carrier family 22 member 3 (SLC22A3). The most downregulated genes (with significant *p*-values) are given in Table 2 and shown in Fig. 2b and also include genes regulating proliferation, cell differentiation, and intracellular signaling such as mitochondrial peptide deformylase (PDF), Suppressor APC domain-containing protein 2 (SAPCD2), Actin filament-associated protein 1-like 2 (AFAP1L2), Leucine-rich repeat-containing protein 17 (LRRC17), Coagulation factor XIII A chain (F13A1), Endothelial cell-specific molecule 1 (ESM1), CMRF35-like molecule 7 (CD300LB), Anoctamin-4 (ANO4), and Transmembrane protein 190 (TMEM190).

**Figure 2.**
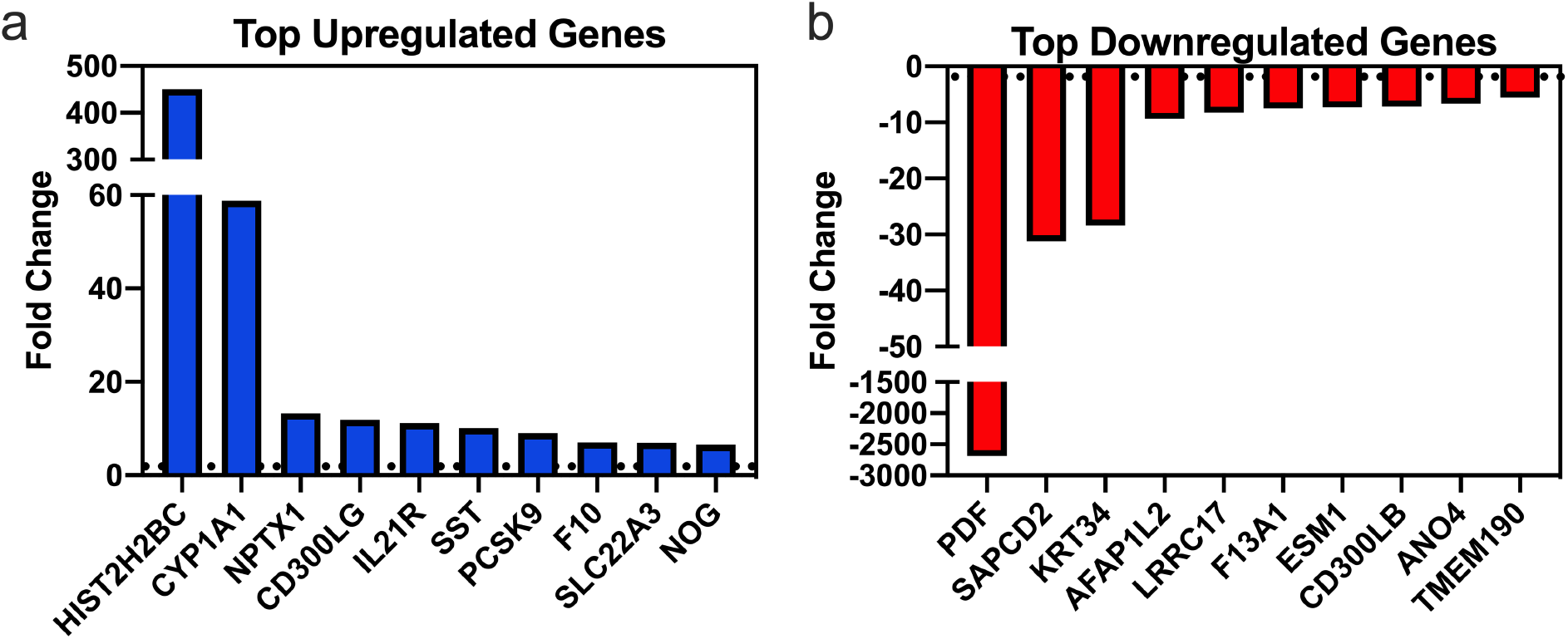
Most upregulated and downregulated genes in NP cells as determined by gene specific analysis. (a) Fold change values of 10 most upregulated genes (LMP gel v. TCPS). (b) Fold change values of 10 most downregulated genes (LMP gel v. TCPS). Horizonal dotted lines denote 2 and −2-fold change, respectively.

**Table 1.** Gene symbol, name, fold change, and *p*-value of 10 most upregulated genes (LMP gels v. TCPS).

| Gene Symbol | Gene Name | Fold Change | *p*-value |
| --- | --- | --- | --- |
| HIST2H2BC | Putative histone H2B type 2-C | 451.39 | 4.5e-2 |
| CYP1A1 | Cytochrome p450 1A1 | 58.79 | 3.6e-5 |
| NPTX1 | Neuronal pentraxin-1 | 13.33 | 4.1e-4 |
| CD300LG | CMRF35-like molecule 9 | 11.93 | 1.9e-2 |
| IL21R | Interleukin-21 receptor | 11.23 | 2.8e-2 |
| SST | Somatostatin | 10.10 | 1.5e-2 |
| PCSK9 | Proprotein convertase subtilisin/kexin type 9 | 8.99 | 3.3e-2 |
| F10 | Coagulation factor X | 7.08 | 4.4e-2 |
| SLC22A3 | Solute carrier family 22 member 3 | 6.97 | 1.1e-2 |
| NOG | Noggin | 6.48 | 1.0e-3 |

**Table 2.** Gene symbol, name, fold change, and *p*-value of 10 most downregulated genes (LMP gels v. TCPS).

| Gene Symbol | Gene Name | Fold Change | *p*-value |
| --- | --- | --- | --- |
| PDF | Peptide deformylase,<br>Mitochondrial | -2683 | 1.6e-4 |
| SAPCD2 | Suppressor APC domain-containing protein 2 | -31.2 | 4.7e-3 |
| KRT34 | Keratin, type I cuticular Ha4 | -28.35 | 3.9e-2 |
| AFAP1L2 | Actin filament-associated protein<br>1-like 2 | -9.36 | 1.7e-2 |
| LRRC17 | Leucine-rich repeat-containing<br>protein 17 | -8.25 | 1.1e-3 |
| F13A1 | Coagulation factor XIII A chain | -7.45 | 1.3e-5 |
| ESM1 | Endothelial cell-specific<br>molecule 1 | -7.25 | 4.0e-2 |
| CD300LB | CMRF35-like molecule 7 | -7.13 | 4.6e-2 |
| ANO4 | Anoctamin-4 | -6.68 | 9.5e-4 |
| TMEM190 | Transmembrane protein 190 | -5.56 | 1.4e-2 |

### 3.2 Markers of NP Cell and Fibroblast Phenotype Are Differentially Regulated by Culture on LMP Gels

When cells were cultured on the LMP gel, several targets previously identified as markers of the NP cell phenotype ^12^ were upregulated including Noggin (NOG) and Integrin a6 (ITGA6) (Fig. 3a). Other markers of the NP phenotype that were moderately increased (but did not exceed a fold change value of 2) included Aggrecan (ACAN), Forkhead Box F1 (FOXF1), Carbonic anhydrase 12 (CA12), and Hypoxia-inducible factor 1 (HIF1A). Markers including Neuropilin 1 (NRP1) and Proteoglycan 4 (PRG4) were downregulated (Fig. 3a) while others were modestly reduced in expression (but did not reach a fold change value of −2) including: Cytokeratin 19 (KRT19), N-cadherin (CDH2), Solute carrier family 2, facilitated glucose transporter member 1 (SLC2A1), Collagen alpha-1(II) chain (COL2A1), and Brain acid soluble protein 1 (BASP1). Expression of a panel of fibroblast markers was also examined ^46–50^ (Fig. 3b). While some genes in the cells cultured on LMP gels were observed to be upregulated (Complement decay-accelerating factor also known as CD55 molecule, CD55, and Endosialin, CD248), others had reduced expression including: CCN family member 2 (CTGF), Vimentin (VIM), Collagen alpha-1 (I) chain (COL1A1), Prolyl endopeptidase FAP (FAP), Actin, aortic smooth muscle (ACTA2), and Vascular cell adhesion protein 1 (VCAM1).

**Figure 3.**
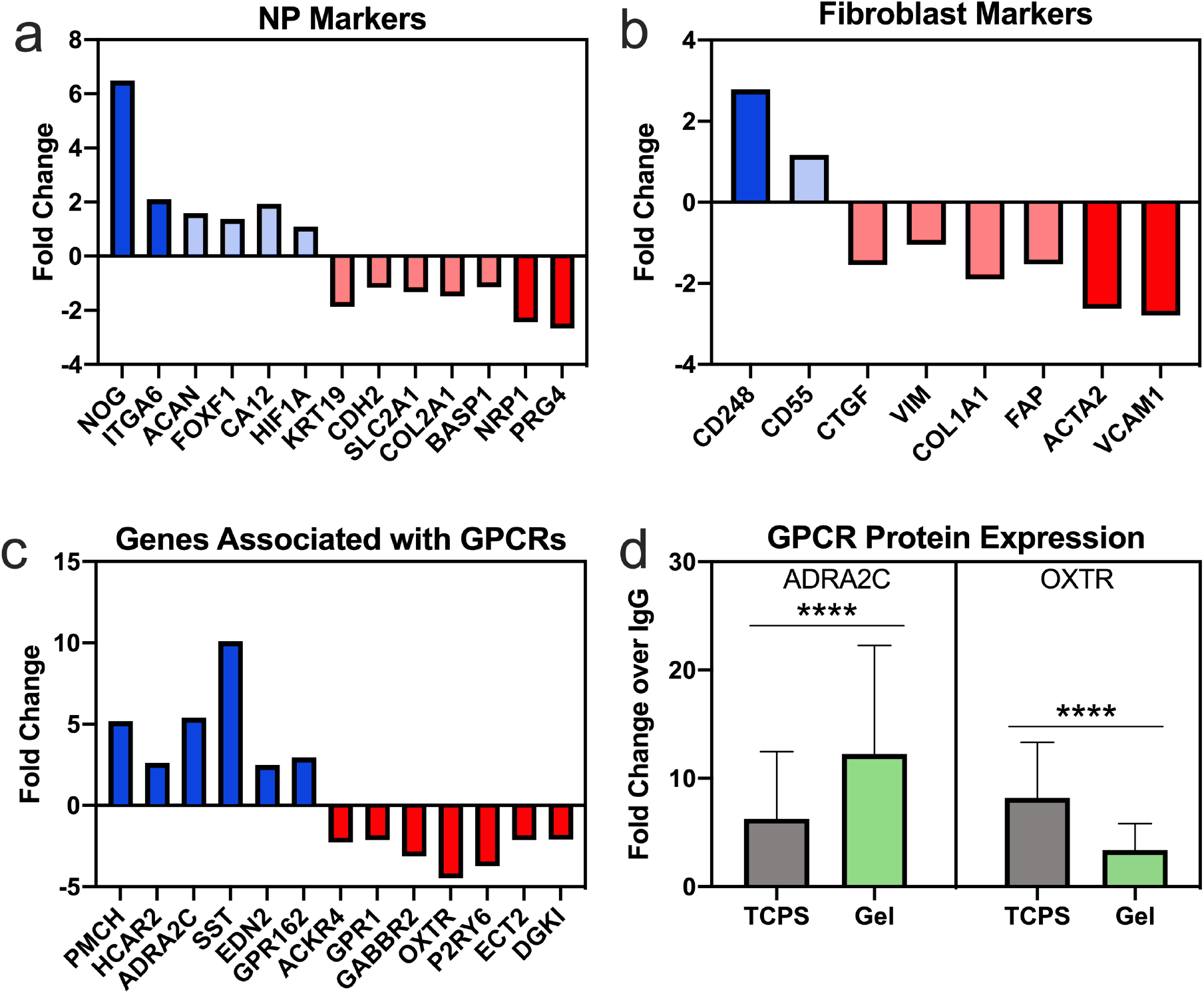
Expression of genes associated with NP and fibroblast cell phenotype of GPCRs. (a) Expression of a panel of NP phenotypic markers, (b) fibroblast markers and (c) up- and downregulated genes associated with G protein coupled receptors (GPCRs). For a-c: Fold change = LMP gel v. TCPS; bars with opaque color had fold change values greater than or equal to 2 or less than or equal to −2). (d) Validation of GPCR protein expression; fold change = expression on TCPS (black bar) or LMP gel (green bar)/expression in the IgG control group. **** *p* < 0.0001.

### 3.3 Genes Related to G Protein-Coupled Receptors Are Among Both Upregulated and Downregulated Genes

Amongst the up- and downregulated genes, 13 genes associated with G protein-coupled receptors (GPCR) were identified (Fig. 3c). These genes were: Pro-MCH (PMCH), Hydroxycarboxylic acid receptor 2 (HCAR2), Alpha-2C adrenergic receptor (ADRA2C), Somatostatin (SST), Endothelin-2 (EDN2), and Probable G-protein coupled receptor 162 (GPR162). Other GPCR-associated genes were downregulated and included: Atypical chemokine receptor 4 (ACKR4), G-protein coupled receptor 1 (GPR1), Gamma-aminobutyric acid type B receptor subunit 2 (GABBR2), Oxytocin receptor (OXTR), P2Y purinoceptor 6 (P2RY6), Protein ECT2 (ECT2), and Diacylglycerol kinase iota (DGKI). The gene expression results were validated at the protein level for ADRA2C and OXTR, representative of up-regulated and down-regulated GPCR-associated genes respectively (Fig. 3d). These two markers were chosen for validation as ADRA2C has been observed to be downregulated in degenerative discs ^51^ and OXTR has been studied in the context of musculoskeletal development, neurogenic pain and inflammation ^52–54^ and have known agonists and antagonists ^55–57^. The protein expression data replicated the results observed through RNA sequencing and confirmed that the differential expression of these two markers was observed at the transcriptomic and proteomic levels.

### 3.4 Gene Set Enrichment Analysis Reveals Pathways and Cellular Components Implicated in NP Cell Culture on LMP Gels

Gene set enrichment was performed on each full list of upregulated or downregulated genes and yielded 721 and 716 significantly enriched gene sets respectively. The enriched gene sets within the upregulated genes represented cellular processes including cellular differentiation, cell signaling, and cytoskeletal regulation (Fig. 4a and Table 3). The top 10 enriched pathways from the upregulated genes include pathways related to organ morphogenesis, including notochord morphogenesis, intracellular signaling pathways (e.g. transforming growth factor beta receptor pathway and bone morphogenic protein (BMP) signaling pathway) and transport of solutes and ions. The gene sets enriched from the downregulated genes included those around themes including cell cycle and DNA processes, extracellular matrix components, and cytoskeletal organization particularly related to cellular division (Fig. 4b and Table 3). Furthermore, the expression of upregulated and downregulated genes implicated in gene sets of particular interest given NP cell biology were examined. In the notochord morphogenesis (GO term: 0048570) gene set, noggin (NOG) and protein WNT-11 (WNT11) were upregulated while this pathway was not associated with any significantly downregulated genes (Fig. 5a). Other gene sets including Cytoskeleton Organization (GO: 0007010), Transforming growth factor beta receptor signaling pathway (GO: 0007179), Signaling (GO: 0023052) were shown to be associated with both upregulated and downregulated genes (Fig. 5b-d).

**Figure 4.**
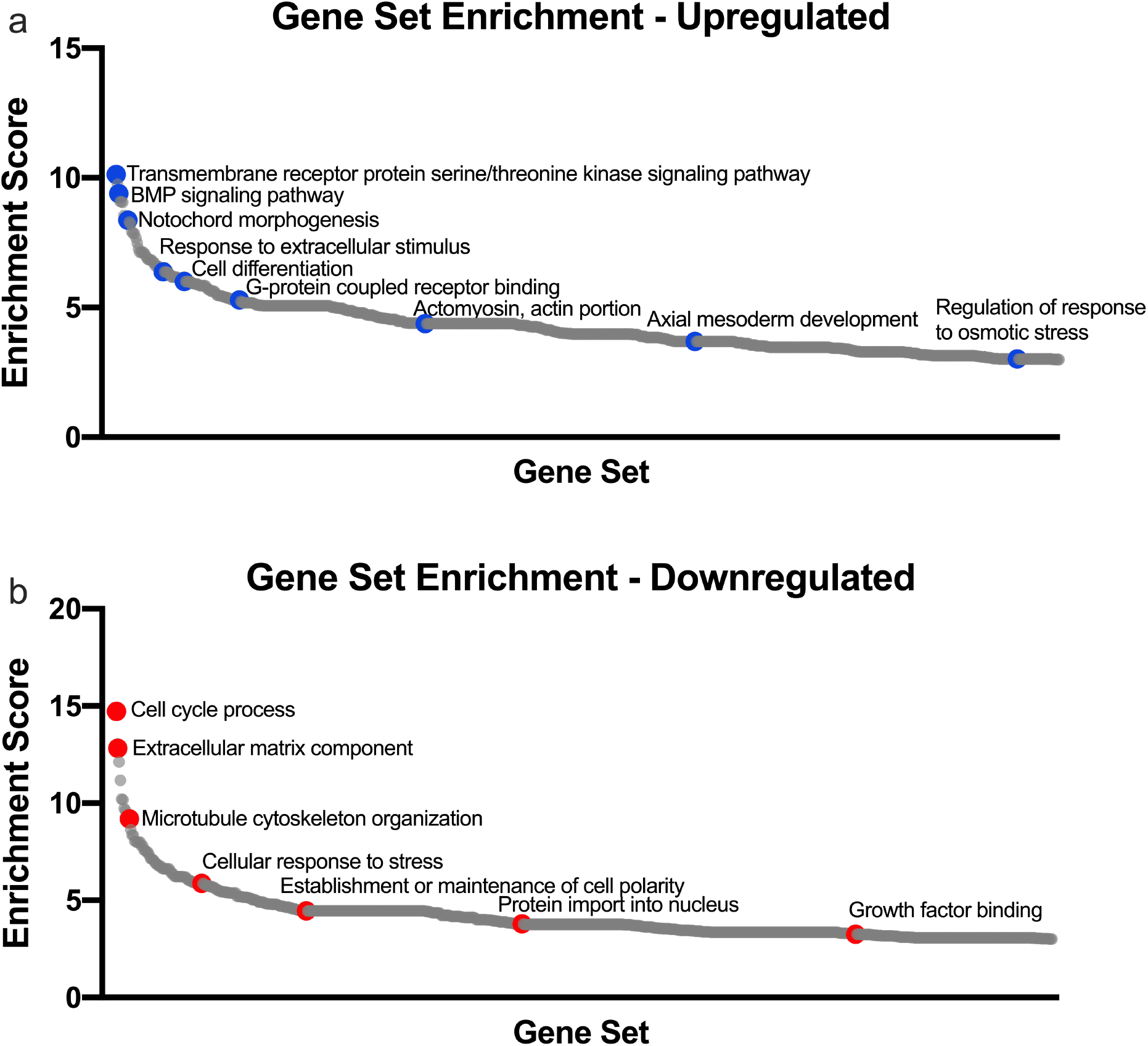
Enriched gene sets in the (a) upregulated and (b) downregulated genes with notable gene sets labeled.

**Table 3.** Top 10 most enriched gene sets associated with the upregulated and downregulated genes.

| Top Gene Sets Enriched – Upregulated gel v. TCPS |  |
| --- | --- |
| 1 | transmembrane receptor protein serine/threonine kinase signaling pathway |
| 2 | antiporter activity |
| 3 | solute:proton antiporter activity |
| 4 | formation of primary germ layer |
| 5 | BMP signaling pathway |
| 6 | zygotic specification of dorsal/ventral axis |
| 7 | ion antiporter activity |
| 8 | transforming growth factor beta receptor signaling pathway |
| 9 | notochord morphogenesis |
| 10 | alpha-2A adrenergic receptor binding |

| Top Gene Sets Enriched – Downregulated gel v. TCPS |  |
| --- | --- |
| 1 | cell cycle process |
| 2 | extracellular matrix component |
| 3 | DNA metabolic process |
| 4 | chromosomal region |
| 5 | mitotic cell cycle process |
| 6 | spindle organization |
| 7 | microtubule cytoskeleton organization involved in mitosis |
| 8 | DNA replication initiation |
| 9 | DNA replication |
| 10 | microtubule cytoskeleton organization |

**Figure 5.**
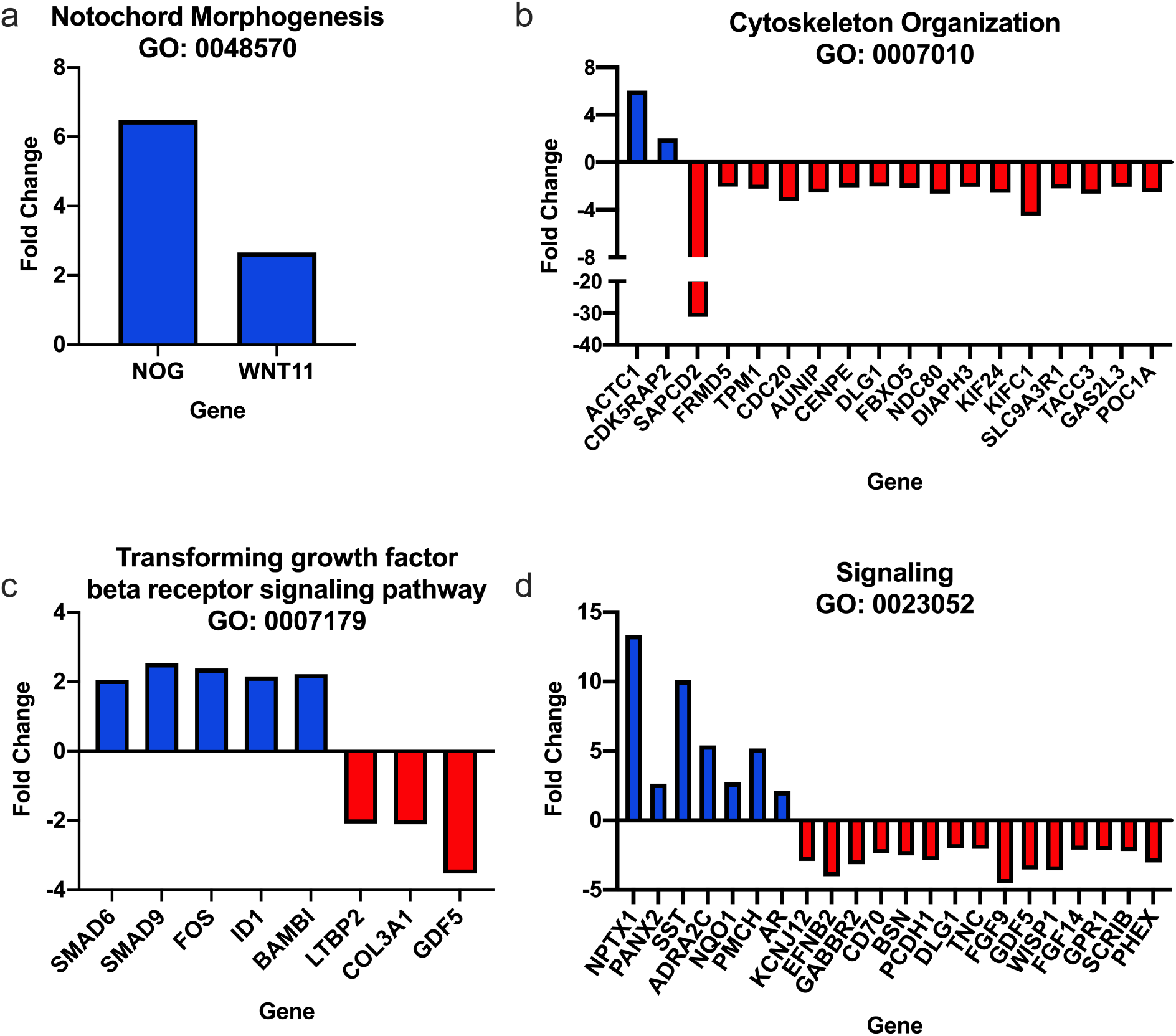
Fold change values (gel v. TCPS) of genes associated with gene ontology terms. (a) Notochord morphogenesis, (b) Cytoskeleton Organization, (c) Transforming growth factor beta receptor signaling pathway, and (d) Signaling.

## 4 Discussion

It has widely been demonstrated that many cell types are able to robustly sense and respond to physical and chemical features of their environment ^20,27,58^. These responses can be directed by using engineered substrates with defined features, such as the LMP gel studied here, to promote cell differentiation or phenotypes *in vitro* ^38,39,59–61^ or for treatment of diseases *in vivo* ^28,62–64^. Characteristics that have been shown to modulate cell behavior in 2D include the stiffness of the underlying substrate, ligand presentation, geometry of adhesive area (as controlled through micropatterning), and ligand density ^9,30,35,39,43,59,65–68^. We have recently demonstrated that a stiff gel (15% PEG) conjugated with equimolar amounts of the laminin-mimetic peptides AG73 and IKVAV could be used to promote degenerative cells to assume a phenotype consistent with healthy, juvenile NP cells ^14^. 15% PEG gels functionalized with 100 µM of peptide (IKVAV and AG73) promoted increased NP cell rounding and decreased actin alignment compared to equally stiff gels functionalized with 400 µM of peptide ^14^. Furthermore, cell shape and cytoskeletal organization on the stiff gels with the lower peptide density replicated behaviors seen in cells cultured on 4% PEGLM ^14^. Additionally, this prior work demonstrated an ability for the peptide-presenting substrates to promote an increase in gene expression of five select NP markers compared to cells cultured upon TCPS ^14^. Rather than only focus on pre-selected molecular targets, in the current study, we sought to examine the broader profile of differential mRNA expression for degenerative NP cells cultured on these substrates. The data presented herein suggest an ability for the biomaterial to alter pathways and cellular function in ways that corroborate, as well as expand on, the previous study.

Results from both hierarchical clustering and PCA analysis demonstrate separation in the transcriptomes of degenerative NP cells cultured on the gel compared to TCPS, a finding that was consistent across samples. Upon further examination, 148 genes were found to be significantly upregulated while 277 genes were identified as significantly downregulated. These findings, like prior literature, suggest that features of the underlying substrate can control global gene expression in cells ^30,60,69^.

The data further demonstrate that culture of degenerative adult human NP cells on gels functionalized with IKVAV and AG73 may cause an increased expression of markers associated with juvenile NP cells, including NOG and ITGA6. Furthermore, expression of many fibroblastic markers including CTGF, ACTA2, VCAM1, and FAP were downregulated. Taken together with previous data, these findings suggest that adult degenerative NP cell phenotype may be modulated to shift towards a more juvenile NP state through culture on this substrate ^14^. Following culture on the LMP gel, however, several NP and fibroblast markers were found to be downregulated and upregulated respectively. This suggests that mechano-biological cues from the gel may regulate cell behaviors in a manner that may be gene/pathway specific and that substrate-directed cues may be potent regulators of cell behaviors, but further control of cell phenotype may require additional factors (e.g. growth factors or pharmacological agents). This is consistent with previous studies which have shown substrate parameters such as the geometry of the adhesive area have been able to promote MSC differentiation towards a particular lineage but alone may be insufficient to promote all cells to commit to a differentiated state ^59,60,65^.

### Most Differentially Expressed Genes

The NP marker NOG which is associated with notochord development was amongst the most highly upregulated genes ^6^. Other highly upregulated terms included genes associated with differentiation including epigenetic regulation (HIST2H2BC) and nervous system development (NPTX1). NPTX1 has been shown to be upregulated in MSCs ^70^ as well as being differentially regulated in the context of development and degeneration of the spinal cord and IVD ^71–73^. NPTX1 was found to be differentially regulated in patients with split cord malformation ^71^. Additionally, studies which have explored the effects of TGFβ on IVD cells found that 8 hours of treatment with 5ng/ml TGFβ1 increased NPTX1 expression in sclerotome cells ^73^, whereas, differentiation of human pluripotent stem cells to NP-like cells via activation of TGFβ signaling (10 ng/ml TGFβ3) reduced NPTX1 expression ^72^. Somatostatin (SST) was also highly upregulated following culture upon the LMP gels which is of interest as the ligand SST has been identified as a potential target for the prevention of IVD cell senescence ^74,75^.

Amongst the most highly downregulated genes, several have been associated with angiogenesis, musculoskeletal development, inflammation, and immune response, pathways of interest as they are implicated in the degenerative cascade in the disc. AFAP1L2 (also known as XB130), one of the most highly downregulated genes in this data set, codes for an adaptor protein that mediates intracellular signal transduction through the PI3K/AKT signaling pathway to regulate proliferation, motility and cell survival as well as having implications in inflammation through Src and IL-8 signaling ^76,77^. LRRC17 is a known negative regulator of RANKL (receptor activator of NF-kappaB ligand) ^78^ that has also been shown to be differentially regulated in the context of both nucleus pulposus and chondrocytes, and was also downregulated by culture on the LMP gel ^72,79^. Likewise, a marker of angiogenesis, ESM1, has been shown to be differentially regulated in this work and in a study that directed the differentiation of human pluripotent stem cells towards notochord- and nucleus pulposus-like populations ^72^. The identification of these up- and down-regulated genes provides insight into the impacts on cell behaviors cultured on the LMP gels and further reveal how the gels may serve to mitigate inflammation and angiogenesis if injected into a degenerative disc space. However, further study is required to confirm these functions. These differentially regulated transcripts, thus, represent targets for further analysis including quantification of protein expression, sub-cellular localization, and elucidation of their specific roles in regulating NP cell phenotypes.

### Genes Associated with G Protein-Coupled Receptors

Other targets examined in the present study were genes associated with G protein-coupled receptors (GPCRs) as these are not only regulators of diverse cellular functions, but also a class of receptors that are commonly druggable targets. The data presented 6 genes that were upregulated and 7 that were downregulated which had GO terms associating them with GPCRs. Interestingly, several of these have been associated previously with the musculoskeletal system and specifically IVD or cartilage. PMCH has not previously been associated with NP phenotype, however studies have shown a role for this GPCR-related gene in early neurogenesis, chondrogenesis, and regeneration as well as in neuropathic and inflammatory pain in a murine animal model ^80–83^. HCAR2 has been implicated in macrophage recruitment and inflammatory responses and protein expression of hydroxycarboxylic acid receptor-2 (HCAR2) has been found in chondrocytes and peripheral nerve among other tissues ^84–86^. Similarly, ADRA2C and SST have been found to be biomarkers in chronic pain for temporomandibular disorder and low back pain ^87–89^. Although a study found low ADRA2C expression in cartilage, a recent report found a decrease in expression of this protein was associated with IVD degeneration ^51,90^. GPR162 (also known as GRCA) has been found in neurons, cartilage, and NP cells ^83,86,91,92^. Though the function of this gene is largely unknown, it has been shown to have a role in cytoskeletal organization and to interact with other

G protein-coupled receptors (STRING database v11) ^93,94^. EDN2 has been found to be involved in the inhibition of vascularization in the retina and other tissues including the IVD; interestingly, it has also been suggested that EDN2 may be a master regulator of notochordal cell gene expression ^95–98^. The upregulation of these genes, some of which have been previously implicated in notochord and healthy features of the NP, provide further data to suggest that culture of degenerative human NP cells upon LMP hydrogels may promote a shift towards expression of a more juvenile-like NP phenotype.

Examination of the GPCR-associated genes which were downregulated in cells cultured on the gels also points to several genes previously shown to have a role in musculoskeletal tissue development or pathology. ACKR4 (also referred to as CCRL1, CCR11, CCX CKR) is a member of the Atypical chemokine receptors family that plays a role in regulating inflammatory responses, yet have also been shown to be present embryonically during NP development and in musculoskeletal diseases such as rheumatoid arthritis ^99–103^. GPR1, which was downregulated in the present study, was found to be upregulated in a study of IVD cells subjected to increased osmolarity ^77^. Prior work in bone has shown that GPR1 deficient mice had decreased bone mass and bone marrow density in addition to increased markers of inflammation, although GPR1’s role in NP and IVD health has not been elucidated ^54,104^. DGKI was also reported to be increased in IVD samples subjected to increased osmolarity and has been implicated in rheumatoid arthritis and osteoarthritis ^77,105–108^. Other downregulated GPCR-associated genes (P2RY6, OXTR, GABBR2, and ECT2) have been studied in the context of neurogenic pain, inflammation, and disc or cartilage tissue development/disease ^52,92,109–116^. However, the role of these genes in regulating cell phenotypes in response to engineered substrates has not previously been elucidated and thus require further validation.

### Gene Set Enrichment Provides Biological Context for Differentially Regulated Genes

Examination of enriched gene ontology terms in the up- and downregulated gene sets reveals not only individual gene changes, but also pathways and cellular components that are potentially altered by culture upon the gels. This analysis confirmed the enrichment of terms associated with notochord morphogenesis and transforming growth factor beta receptor signaling pathway, a signaling cascade critical for the formation of the notochord and NP ^73^. Additionally, the BMP signaling pathway was found to be enriched in the upregulated genes. While this pathway has been associated with the development of bone in some cell types (osteoblasts, chondrocytes, and osteoclasts) ^117^, studies have shown that BMP-2, −7, and −12, have been associated with promoting proteoglycan biosynthesis in NP cells ^18,118,119^. Similarly, pathways related to cell differentiation and response to extracellular stimuli were highly enriched in the data set. Pathways enriched in the downregulated genes suggest that following interactions with the gels, genes associated with cell division are reduced and cytoskeletal organization is altered.

Taken together, the results of this study demonstrate that the cell-matrix interactions between NP cells and hydrogels functionalized with IKVAV and AG73 are able to modulate genes and pathways associated with diverse cellular functions including cell phenotype. As cell clustering has previously been observed on the LMP gel ^14^, additional studies are needed to determine the relative roles of the cell-cell and cell-matrix interactions and to determine the impact of individual substrate parameters such as bulk stiffness and ligand presentation (syndecan and integrin-regulated effects) on the modulation of NP cell phenotype. While this study made use of a single-time point, future analysis of the transcriptome at several time points (ex. 2 h, 24 h, and 4 d) could be used identify transient cellular responses including both early and sustained effects of NP cell interactions with the hydrogel. Furthermore, these transient responses could provide insights into the mechanical memory of NP cells, a factor that has been demonstrated to be an important in regulating biosynthesis and phenotype for other cell types ^120,121^ but has largely not been explored in IVD cells.

In summary, data from the study of the RNA transcriptome reveals an upregulation of individual notochordal markers and pathways associated with the notochord and cell differentiation in degenerate human NP cells upon interacting with a peptide-presenting PEG hydrogel. Furthermore, the use of unbiased analysis to explore novel targets has led to the identification of GPCR-associated genes that point to pathways and druggable targets of interest in IVD development and disease. As with all transcriptomic analysis, additional studies are necessary for validating the expression and subcellular localization of the protein products of these genes. The present study is intended to provide insight into the global gene expression changes in degenerative cells cultured upon an engineered substrate, and the role of the substrate in promoting the assumption of a healthy phenotype.

## 5. Acknowledgements

This work was supported by the National Science Foundation (Graduate Research Fellowship Program under Grant No. DGE-1745038), National Institutes of Health (R01AR069588, R01AR077678) and the Spencer T. and Ann W. Olin Fellowship for Women in Graduate Study. The authors would additionally like to thank the Genome Technology Access Center at Washington University in St. Louis for their assistance with the RNA sequencing. The GTAC is partially supported by the NCI Cancer Center Support Grant Number P30 CA91842 and by ICTS/CTSA Grant Number UL1TR002345 from the NCRR, a component of the NIH, and NIH Roadmap for Medical Research. Any opinions, findings, conclusions, or recommendations expressed in this material are those of the authors and do not necessarily reflect the views of these funding sources.

## 6. Author Contributions

M.N.B and J.E.S designed and performed research, analyzed data, and wrote the paper; L.J. helped to perform research; J.M.B, M.P.K, and M.C.G. contributed to data collection. L.A.S contributed to the study conception and design, writing of the manuscript and data interpretation. All authors were involved in revisions to the manuscript and approval of the final version for submission.

